# Sleep selectively stabilizes contextual aspects of negative memories

**DOI:** 10.1101/370965

**Authors:** Roy Cox, Marthe LV van Bronkhorst, Mollie Bayda, Herron Gomillion, Eileen Cho, Elaine Parr, Olivia P Manickas-Hill, Anna C Schapiro, Robert Stickgold

## Abstract

Sleep and emotion are both powerful modulators of the long-term stability of episodic memories, but precisely how these factors interact remains unresolved. We assessed changes in item recognition, contextual memory, and affective tone for negative and neutral memories across a 12 h interval containing sleep or wakefulness in 71 human volunteers. Our data indicate a sleep-dependent stabilization of negative contextual memories, in a way not seen for neutral memories, item recognition, or across wakefulness. Furthermore, retention of contextual memories was positively associated with time spent in non-rapid eye movement sleep. Finally, our results offer partial support for the hypothesis that sleep attenuates emotional responses to previously memorized material.

## Introduction

Episodic memories are rich, multisensory accounts of past events that guide our future behavior. Besides linking together various aspects of the external world into an integrated whole (e.g., objects, locations, temporal order), episodic memories also contain the emotional tone associated with an experience, resulting in qualitative and quantitative differences between emotional and neutral memories ^1^. In particular, negative emotional memories are typically more durable than neutral memories, especially after longer retention intervals ^2–5^.

Sleep plays a fundamental role in processing episodic memories, both stabilizing recent experiences and integrating them with stored knowledge ^6,7^. These processes are thought to rely on the reinstatement of learning-related activity patterns in the neural structures originally encoding the memory, allowing hippocampal memory traces to be recoded in a more stable neocortical format ^8–10^. Sleep-dependent memory consolidation effects are generally more reliable for associative and contextual memories compared to individual item recognition (e.g., ^11,12^), consistent with stronger initial hippocampal involvement for memories with an associative component ^13^.

Some evidence suggests that sleep and emotion interact, with enhanced retention of negative relative to neutral material following sleep ^14–17^, although other studies have failed to find such an effect ^18–22^. Furthermore, while several examinations into the contributions of specific sleep stages to emotional memory consolidation have emphasized the importance of rapid-eye movement (REM) sleep ^17, 23–25^, others point to a role for non-rapid eye movement (NREM) sleep ^26–28^, or find no clear stage dependence ^20, 29–31^. These inconsistencies may be related to various experimental design factors (e.g., sleep vs. wake; full night vs. nap, split-night; sleep-deprivation; targeted memory reactivation), as well as the memory paradigm employed. Specifically, the clearer effect of sleep on hippocampus-dependent associative memories compared to item recognition could result in greater emotional benefits for memories with an associative component, but again, the available evidence is contradictory ^25, 32–34^.

Another factor in sleep-related emotional processing concerns not the memories *per se*, but the modulation of their affective component. Whereas some physiological evidence suggests that sleep serves to decrease emotional reactivity to previously seen emotional items ^35,36^, subjective ratings of emotionality have yielded discrepant findings. Some studies report no difference in emotional reactivity to emotional versus neutral items, with similar decreases ^24^ or increases ^18^ across sleep versus wake, or comparable ratings after NREM and REM sleep for both emotional categories ^23^. Other investigators have described a REM-dependent preservation of negative ratings relative to wake ^19^ or REM deprivation ^37^, while yet others have reported enhanced emotional reactivity after REM-rich sleep ^32,38^.

In sum, sleep’s role in emotional processing remains unclear, both with regards to memory consolidation and affective processing (see ^39^ for a recent review). Here, we re-examine the role of sleep in emotional processing, focusing on both recognition and contextual memory, as well as subjective affective ratings. Participants encoded negative and neutral item-context associations in either the evening or morning, and were tested both immediately after encoding and after a 12 h interval containing sleep or wake. Polysomnography was recorded for the sleep group in order to investigate associations with sleep architecture. We hypothesized that sleep would preferentially stabilize the contextual aspects of negative memories. Given the conflicting findings reviewed above, we did not have strong predictions for recognition memory or affective ratings, nor did we have clear expectations on the involvement of specific sleep stages.

## Results

### Protocol

Seventy-one healthy volunteers underwent a protocol consisting of memory encoding and both immediate and delayed retrieval, with retrieval tests organized around a ~12 h retention interval containing wakefulness or sleep (Fig. 1A). Participants in the wake group (N=25) performed encoding and immediate testing (Test 1) in the morning, and delayed testing at night (Test 2). This timing was reversed for participants in the sleep group (N=46). Furthermore, 62-channel EEG was recorded throughout the procedure in the sleep group to allow sleep staging as well as future examinations of the neural correlates of encoding, retrieval, and memory reactivation during sleep.

**Figure 1.**
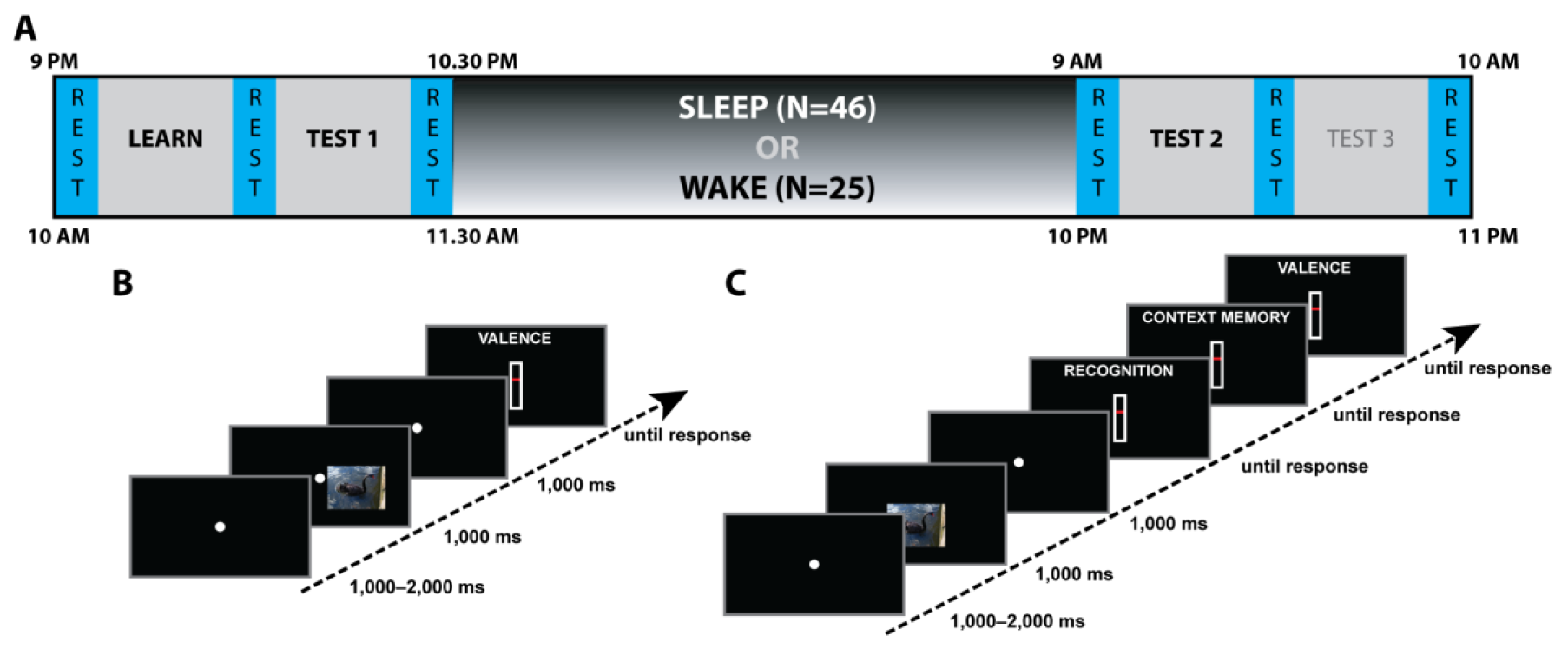
Protocol overview. (A) Study timeline. (B) Trial structure at encoding (with lateralized stimulus presentation). (C) Trial structure at retrieval (with central stimulus presentation).

During encoding, participants maintained fixation in the center of the screen while a series of visual stimuli was briefly presented in either the left or right visual field (VF) (Fig. 1B). Importantly, all pictures shown on one side were of mild positive emotional valence and low arousal (“neutral valence”), while those presented on the opposite side were of strong negative valence and high arousal (“negative valence”); this manipulation served to facilitate future decoding of category-specific EEG signatures (see Discussion). Subjects rated each item’s perceived valence immediately following its presentation on a continuous scale from very negative to very positive. Participants were not told that their memory would be tested later. Assignment of emotional category to VF was counterbalanced across subjects. This additional factor SIDE (negative-left/neutral-right: N=36; negative-right/neutral-left: N=35) was included to ensure that any differences between negative and neutral categories could not be due to asymmetric emotional processing by the two hemispheres ^40–42^.

During the immediate and delayed retrieval tests, subjects viewed previously seen (“old”) and novel (“new”) stimuli presented in the center of the screen (as opposed to lateralized; Fig. 1C). Different old and new items were used for immediate and delayed tests. Subjects then used sequentially presented continuous scales to rate each item’s 1) old/new status (recognition memory), 2) VF of presentation during encoding (contextual memory), and 3) perceived valence. For items rated “new”, subjects rated VF as if they were actually “old” items (see Methods).

### Control Analyses

We first assessed whether time of day affected vigilance and whether that, or the VF of negative item presentation, had any effect on valence ratings or sleep architecture.

#### Vigilance

We determined subjective vigilance in two ways both prior to encoding and prior to the delayed test. Sleepiness, as assessed by the Stanford Sleepiness Scale ^43^ (SSS), did not differ between the sleep and wake groups either before encoding (sleep: 2.6 ± 0.8; wake: 2.2 ± 0.6; Wilcoxon Rank Sum: Z=1.7, P=0.09) or after the 12 h interval (sleep: 2.5 ± 1.1; wake: 2.4 ± 0.9; Z=0.2, P=0.81). Similarly, self-reported ability to concentrate (continuous scale from 0–100) did not differ systematically pre-encoding (sleep: 74.4 ± 15.6; wake: 80.6 ± 13.8; t(69)=1.7, P=0.10) or before the delayed test (sleep: 74.1 ± 18.4; wake: 75.8 ± 15.9; t(69)=0.4, P=0.70), indicating no time of day effect on vigilance.

#### Encoding Valence

During encoding, subjects rated the emotional valence of each picture on a scale from –1 (negative) to +1 (positive) immediately following its lateralized presentation. An ANOVA with between-subject factor GROUP (sleep/wake) and within-subject factor VALENCE (negative/neutral) on subjects’ trial-averaged ratings indicated that negative items were rated significantly more negatively than neutral ones (main effect of VALENCE), with no effect of GROUP or GROUP*VALENCE interaction (Table 1–*encoding*). Post-hoc tests indicated that the valence effect was present in both groups (sleep: t(45)=30.4, P<10^−30^; wake: t(24)=16.0, P<10^−13^), whereas subjective ratings did not differ systematically between the sleep and wake groups for either neutral (t(69)=1.7, P=0.09) or negative (t(69)=0.8, P=0.41) items, indicating no time of day effect on perceived valence. Moreover, single-subject independent t tests comparing negative and neutral items indicated that each of the 71 individuals rated negative items significantly more negatively than neutral items (all P_adj_<10^-14^, where P_adj_ indicates the adjusted P value using the False Discovery Rate [FDR] ^44^). We further added SIDE (negative-left/negative-right) as a between-subject factor, but found no significant main or interaction effects involving this factor (all F(1,67)<2.5, all P>0.12), indicating that the VF where each emotional category was presented did not impact ratings. Thus, these results establish that subjects can accurately gauge the emotional valence of stimuli presented away from fixation, regardless of presentation side, and that these effects are similar in the morning and evening.

**Table 1.**
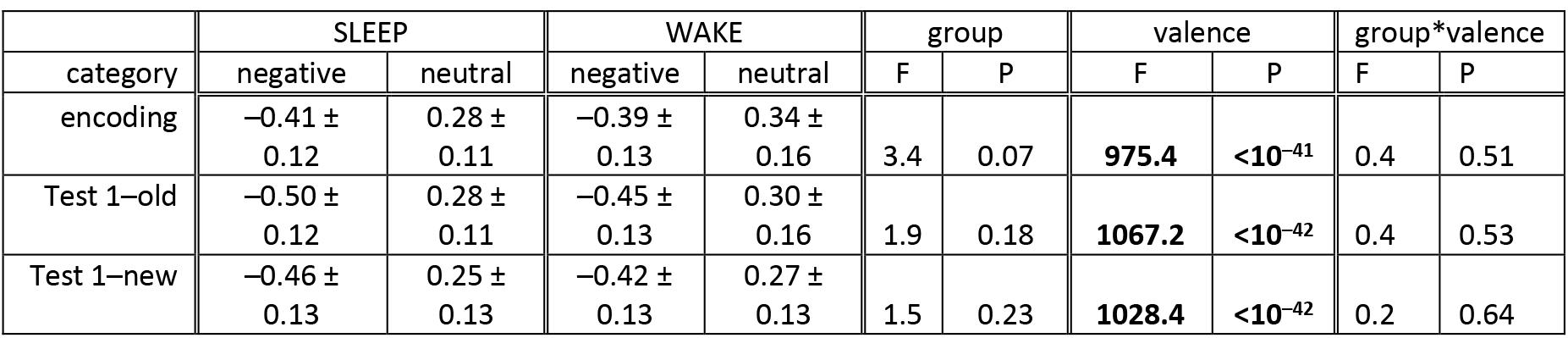
Valence ratings at encoding and immediate test (mean ± SD). F test degrees of freedom: (1,69). Significant effects indicated in bold.

| category | SLEEP |  | WAKE |  | group |  | valence |  | group*valence |  |
| --- | --- | --- | --- | --- | --- | --- | --- | --- | --- | --- |
|  | negative | neutral | negative | neutral | F | P | F | P | F | P |
| encoding | -0.41 ± 0.12 | 0.28 ± 0.11 | -0.39 ± 0.13 | 0.34 ± 0.16 | 3.4 | 0.07 | **975.4** | **<10<sup>-41</sup>** | 0.4 | 0.51 |
| Test 1–old | -0.50 ± 0.12 | 0.28 ± 0.11 | -0.45 ± 0.13 | 0.30 ± 0.16 | 1.9 | 0.18 | **1067.2** | **<10<sup>-42</sup>** | 0.4 | 0.53 |
| Test 1–new | -0.46 ± 0.13 | 0.25 ± 0.13 | -0.42 ± 0.13 | 0.27 ± 0.13 | 1.5 | 0.23 | **1028.4** | **<10<sup>-42</sup>** | 0.2 | 0.64 |

#### Sleep Architecture

Overall sleep architecture for the 46 overnight participants was in line with typical values for young, healthy subjects (Table 2). We compared sleep parameters as a function of SIDE to examine if the VF where negative vs. neutral items had been presented affected subsequent sleep. Time and percentages spent in each sleep stage did not differ between groups. We did observe differences significant at an uncorrected threshold in parameters related to sleep quality (total sleep time, sleep efficiency, sleep latency), which were driven by two outliers in the negative-left group with unusually long sleep latencies (76 and 81.5 min; all others <36 min), one of whom also had the lowest observed sleep efficiency (72.6%). However, no comparison survived correction for multiple testing. Thus, macroscopic sleep structure did not vary depending on the hemisphere initially processing negative or neutral information prior to sleep.

**Table 2.** Sleep architecture parameters (mean ± SD). WASO: wake after sleep onset; P_adj_: adjusted P value using FDR. Entries in bold indicate significance (P<0.05).

|  | overall | negative-left | negative-right | statistics |  |  |
| --- | --- | --- | --- | --- | --- | --- |
|  | N=46 | N=22 | N=24 | t(44) | P | P<sub>adj</sub> |
| N1 (%) | 3.9 ± 2.7 | 4.4 ± 2.8 | 3.4 ± 2.6 | 1.3 | 0.20 | 0.35 |
| N2 (%) | 51.1 ± 6.2 | 52.0 ± 6.1 | 50.3 ± 6.3 | 0.9 | 0.37 | 0.46 |
| N3 (%) | 24.5 ± 5.7 | 24.0 ± 5.8 | 25.1 ± 5.7 | -0.7 | 0.51 | 0.59 |
| REM (%) | 20.5 ± 4.3 | 19.7 ± 4.2 | 21.3 ± 4.3 | -1.3 | 0.22 | 0.35 |
| N2-N3 (%) | 75.6 ± 4.7 | 75.9 ± 4.0 | 75.4 ± 5.3 | 0.4 | 0.70 | 0.73 |
| N1-N2-N3 (%) | 79.5 ± 4.3 | 80.3 ± 4.2 | 78.7 ± 4.3 | 1.3 | 0.22 | 0.35 |
| N1 (min) | 18.0 ± 12.1 | 20.1 ± 12.1 | 16.1 ± 12.0 | 1.1 | 0.27 | 0.36 |
| N2 (min) | 243.0 ± 34.3 | 241.1 ± 34.5 | 244.7 ± 34.8 | -0.3 | 0.73 | 0.73 |
| N3 (min) | 116.7 ± 28.4 | 111.0 ± 27.5 | 122.0 ± 28.7 | -1.3 | 0.19 | 0.35 |
| REM (min) | 98.0 ± 23.6 | 91.9 ± 24.1 | 103.6 ± 22.2 | -1.7 | 0.10 | 0.35 |
| N2-N3 (min) | 359.7 ± 34.9 | 352.1 ± 33.1 | 366.7 ± 35.7 | -1.4 | 0.16 | 0.35 |
| N1-N2-N3 (min) | 377.7 ± 30.8 | 372.2 ± 31.4 | 382.8 ± 30.0 | -1.2 | 0.25 | 0.36 |
| WASO (min) | 22.6 ± 24.6 | 27.6 ± 27.2 | 18.0 ± 21.6 | 1.3 | 0.19 | 0.35 |
| total sleep (min) | 475.7 ± 36.9 | 464.1 ± 39.7 | 486.4 ± 31.4 | -2.1 | **0.04** | 0.21 |
| sleep latency (min) | 14.9 ± 16.7 | 20.6 ± 21.5 | 9.6 ± 8.0 | 2.4 | **0.02** | 0.18 |
| sleep efficiency (%) | 89.4 ± 5.6 | 87.2 ± 6.2 | 91.5 ± 4.1 | -2.8 | **0.008** | 0.13 |

### Contextual Memory

We next analyzed subjects’ memory for the VF in which each item had been presented during encoding (i.e., contextual memory). Continuous left/right ratings were dichotomized and relabeled “correct” and “incorrect” according to original presentation side. In keeping with previous aproaches ^13,45^, we focus below on contextual memory for correctly recognized items (i.e., hits).

#### Immediate Test

At immediate testing, subjects correctly indicated the original presentation side significantly above chance in each condition (negative-sleep: 73.9 ± 14.0%; neutral-sleep: 74.7 ± 16.9%; negative-wake: 75.6 ± 16.5%; neutral-wake: 78.3 ± 15.5%; t tests vs. 50%: all P<10^−7^). An ANOVA with between-subject factor GROUP and within-subject factor VALENCE did not reveal significant baseline differences (all F(1,69)<0.8, all P>0.37). Additional analyses including factor SIDE are presented in Supplementary Information.

#### Change over 12 h

Next, we evaluated the change in contextual memory across the 12 h interval (delayed –immediate). As predicted, we observed a selective preservation of contextual memory across 12 h for negative items in the sleep group (Fig. 2A). Whereas performance decreased across wake for negative (–11.1 ± 13.8%; one sample t test vs. zero: t(24)=4.0, P=0.0005) and neutral items (–5.5 ± 15.2%; t(24)=1.8, P=0.08), and for neutral items across sleep (–7.8 ± 14.2%; t(45)=3.7, P=0.0005), memory for negative items was unchanged across sleep (0.2 ± 14.3%; t(45)=0.1, P=0.94). Across conditions, there was a significant GROUP*VALENCE interaction (F(1,69)=5.7, P=0.02), a main effect of GROUP (F(1,69)=4.2, P=0.04), but no effect of VALENCE (F(1,69)=0.2, P=0.67). Post hoc tests indicated that the change in contextual memory for negative items in the sleep group was significantly different from that for neutral items in the same group (paired t(45)=2.4, P=0.02), and from that for negative items in the wake group (independent t(69)=3.2, P=0.002). The comparison with the neutral-wake condition was in the same direction as for the other two comparisons, but did not reach significance (t(69=1.6, P=0.12). Within the wake group, forgetting of negative and neutral items did not differ significantly (t(24)=–1.5, P=0.26). Further analyses indicated that these effects did not depend importantly on whether negative items had been presented to the left or right during encoding (Supplementary Information).

**Figure 2.**
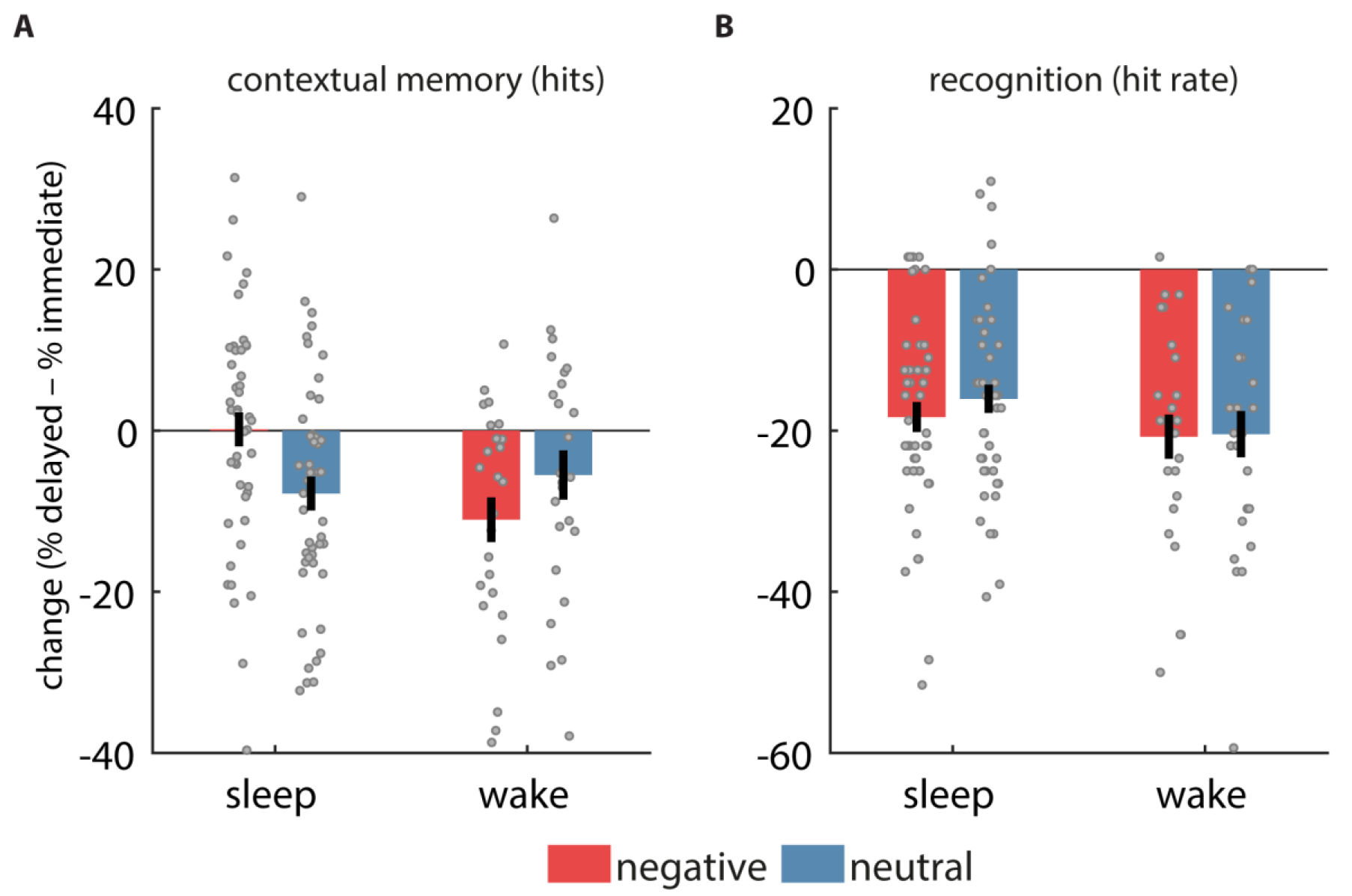
Change in memory across 12 h. (A) Contextual memory for hits was selectively preserved for negative items in the sleep group. (B) Recognition memory (hit rate) dropped similarly for each condition. Error bars reflect standard error of the mean with between-subject variability removed.

#### Contextual Memory Control Analyses

An exit questionnaire indicated that 77% of subjects (total: 55/71; sleep: 36/46; wake: 19/25) became explicitly aware of the mapping between VF and emotional category at some point during the protocol. Hence, it is possible that, during retrieval, participants determined each item’s original presentation side solely based on its perceived valence. To address this issue, we examined baseline performance at immediate test in two sets of analyses. Because sleep and wake groups did not differ at immediate test with respect to either contextual memory or valence, groups were combined.

First, we performed within-subject across-trial correlations, where we correlated items’ valence ratings with their corresponding continuous “correct side/incorrect side” confidence ratings. This was done separately for negative and neutral, and old and new, items. We then extracted the slopes of these regression lines to examine the link between item valence and spatial judgment at the group level. Slopes differed significantly from zero for all four item categories (all t(70)>5.4, all P<10^−6^), indicating that more extreme valence ratings were associated with higher-confidence scores towards the correct side. We reasoned that if subjects exclusively relied on item valence to determine item placement, these relations should be similar for old and new items.

However, slopes differed significantly between old and new items for both valence categories (negative-old: −0.42 ± 0.32; negative-new: −0.30 ± 0.37; t(70)=3.8, P=0.0003; neutral-old: 0.33 ± 0.31; neutral-new: 0.23 ± 0.35; t(70)=2.6, P=0.01), indicating that side judgments for old items went beyond the “baseline” association with valence seen for new items.

Second, if subjects merely relied on item valence, contextual memory for hits should not differ from performance for items with other recognition statuses (i.e., correct rejections, misses, and false alarms), as valence information is equally available in each of these cases. However, when we compared baseline VF ratings for hits to the other three categories, separately for negative and neutral items, we found significantly higher performance for hits for 5/6 comparisons (paired t tests, all P_adj_<0.00004) except for hits vs. false alarms for negative items (P_adj_=0.70). To follow up on the latter finding, we examined the change in performance on the contextual task for false alarms across 12 h. Unlike the differential retention of encoding side for previously presented negative and neutral items across sleep, no such valence-related difference was seen for proportions of false alarms assigned to the “correct” side (negative: –1.5 ± 22.9%; neutral: −7.5 ± 32.9%; t(43)=1.0, P=0.34), suggesting that the emotional advantage was tied to previously encoded items.

Combined, these findings suggest that while valence-based strategies contributed to judgments about encoding side, contextual memory for hits relied on additional memory-related processes. Moreover, the empirical finding that sleep, but not wake, selectively stabilizes contextual memories of negative, but not neutral, items cannot be explained by a pure valence-based account.

#### Contextual Memory and Sleep Parameters

Next, we examined whether overnight changes in contextual memory for hits were related to sleep parameters (all variables in Table 1, except WASO, sleep efficiency, and sleep latency). Separate analyses for negative, neutral, and pooled emotional categories, indicated a strong positive correlation between percentage of time spent in NREM (N1+N2+N3) sleep and contextual memory change across emotional categories (Fig. 3; Spearman R=0.33, Robust regression P=0.003). A corresponding negative correlation was found for REM percentage. These correlations remained significant after correction for multiple comparisons (P_adj_=0.05). We also observed a negative correlation with minutes spent in REM (R=–0.33, P=0.01), although this relation did not survive correction for multiple comparisons (P_adj_=0.16). Correlations with individual emotional categories were not significant for any sleep parameter (all P_adj_>0.40), nor were correlations between time spent in NREM sleep and contextual memory change different for the two emotional categories (z test for correlated correlations: Z=0.34, P=0.63).

**Figure 3.**
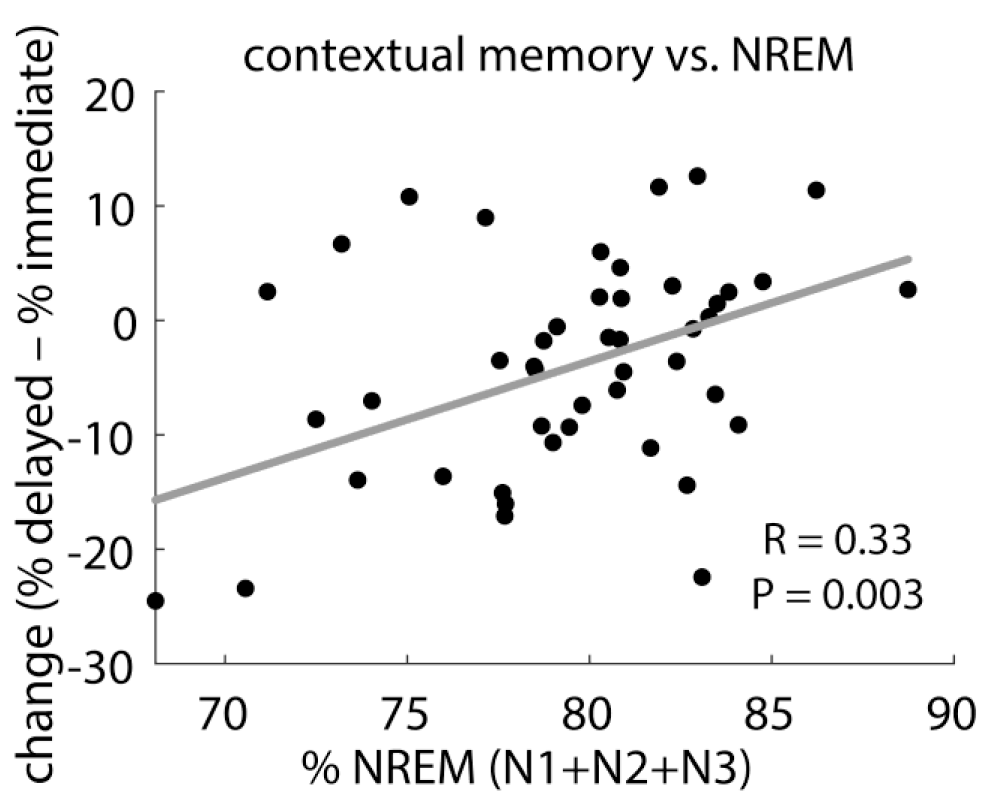
Overnight change in contextual memory for hits, pooled across emotional categories, was positively related to NREM sleep percentage. P value (uncorrected) and regression line from robust fit, R value from Spearman correlation.

In sum, larger proportions of NREM (and smaller proportions of REM) sleep are related to better retention of contextual details regardless of emotional category, suggesting that NREM sleep promotes consolidation of episodic memories in a valence-independent fashion.

### Item Recognition

Next, we assessed old/new recognition memory by dichotomizing subjects’ continuous old/new response to each stimulus as “old” or “new”, and then calculating standard metrics of hit rate (HR), and discriminability (d’). Performance at immediate test for these metrics, as well as for the false alarm rate (FAR), is reported in Supplementary Information.

#### Change over 12 h

We evaluated changes in recognition memory across the 12 h interval (delayed – immediate) as a function of GROUP and VALENCE (Fig. 2B). HR decreased significantly for both negative and neutral stimuli in both groups (one-sample t tests vs. zero: all P<10^−6^), indicating robust forgetting. However, neither GROUP nor VALENCE affected the rate of forgetting (statistics in Table 3–*HR*). Adding SIDE as a between-subject factor did not reveal additional significant main or interaction effects involving this factor (all F(1,67)<2.5, all P>0.12), indicating that forgetting was similar when negative items had originally been presented in the left or right VF. Similar to findings for HR, while subjects’ ability to discriminate old from new items decreased significantly in each condition (one-sample t tests vs. zero: all P<10^−4^), forgetting was not affected by GROUP or VALENCE (all F(1,69)<1.9, all P>0.17; Table 3–*d’*). Again, adding SIDE did not yield significant effects (all F(1,67)<2.7, all P>0.10). Changes in FAR are reported in Supplementary Information.

**Table 3.**
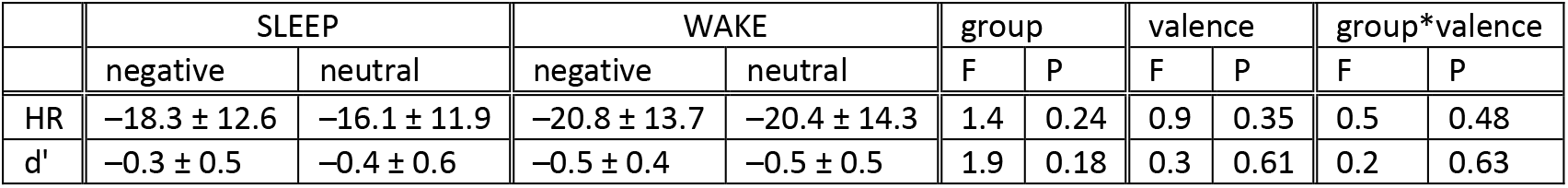
Change in recognition memory across 12 h (mean ± SD). F test degrees of freedom: (1,69).

|  | SLEEP |  | WAKE |  | group |  | valence |  | group*valence |  |
| --- | --- | --- | --- | --- | --- | --- | --- | --- | --- | --- |
|  | negative | neutral | negative | neutral | F | P | F | P | F | P |
| HR | $-18.3 \pm 12.6$ | $-16.1 \pm 11.9$ | $-20.8 \pm 13.7$ | $-20.4 \pm 14.3$ | 1.4 | 0.24 | 0.9 | 0.35 | 0.5 | 0.48 |
| d' | $-0.3 \pm 0.5$ | $-0.4 \pm 0.6$ | $-0.5 \pm 0.4$ | $-0.5 \pm 0.5$ | 1.9 | 0.18 | 0.3 | 0.61 | 0.2 | 0.63 |

For the sleep group we examined whether sleep parameters correlated with overnight memory changes. For each recognition metric (HR, FAR, d’), we performed separate analyses for negative, neutral, and pooled emotional categories. We found no significant correlations (FDR correction across 39 comparisons for each recognition metric: all P_adj_>0.58).

In sum, while overall recognition memory deteriorated markedly across a 12 h interval, wake vs. sleep did not affect the overall rate of forgetting, nor did it impact retention of negative and neutral items differently, contrasting markedly with the effects reported for contextual memory.

### Emotional Valence

Next, we examined whether and how sleep influences subjective ratings of item valence from immediate to delayed testing. We assessed valence separately for old and new pictures to determine whether changes in emotional reactivity are specific to memorized items.

#### Immediate Test

We first sought to ensure that baseline valence ratings at immediate test differed reliably between negative and neutral items, and did so similarly for the sleep and wake groups. Separate ANOVAs for old and new items indicated highly significant main effects of VALENCE, with no effect of GROUP and no GROUP*VALENCE interaction, indicating no effect of time of day on emotional ratings (Table 1, *Test 1–old* and *Test 2– new*). Post hoc paired t tests indicated that negative and neutral ratings differed for each combination of sleep/wake and old/new (all P<10^−13^), and for every individual (all P_adj_<10^−5^). VF of presentation at encoding (SIDE) did not impact affective ratings at immediate test, for either old or new items (all F(1,67)<1.7, all P>0.20).

Adding the factor OLD/NEW to the VALENCE*GROUP ANOVA again revealed a significant effect of VALENCE (F(1,69)=1126.6, P<10^−43^), as well as an OLD/NEW*VALENCE interaction (F(1,69)=28.0, P<10^−5^), but no effects involving GROUP (all other F(1,69)<1.8, P>0.18). Post hoc tests revealed that the OLD/NEW*VALENCE interaction stemmed from more negative ratings to negative items for old compared to new items (sleep: t(45)=3.3, P=0.002; wake: t(24)=2.3, P=0.03; Table 1), and from old neutral items being rated more positively than new neutral items in both groups (sleep: t(45)=3.2, P=0.002; wake: t(24)=2.3, P=0.03; Table 1). These findings indicate that items previously encountered during encoding elicit a stronger emotional response at test than novel items, with this emotional potentiation effect similar for negative and neutral items and independent of time of day.

#### Change over 12 h

We next analyzed changes in valence ratings across the 12 h interval (delayed – immediate; Fig. 4). An ANOVA with factors OLD/NEW, GROUP, and VALENCE yielded significant OLD/NEW*VALENCE (F(1,69)=11.9, P=0.001), GROUP*VALENCE (F(1,69)=4.5, P=0.04), and VALENCE (F(1,69)=8.8, P=0.004) effects (all other F(1,69)<1.3, P>0.26). Post hoc tests indicated that whereas a period of wake did not result in significant changes for any condition (one sample t tests vs. zero, all P>0.16), an interval of sleep did.

**Figure 4.**
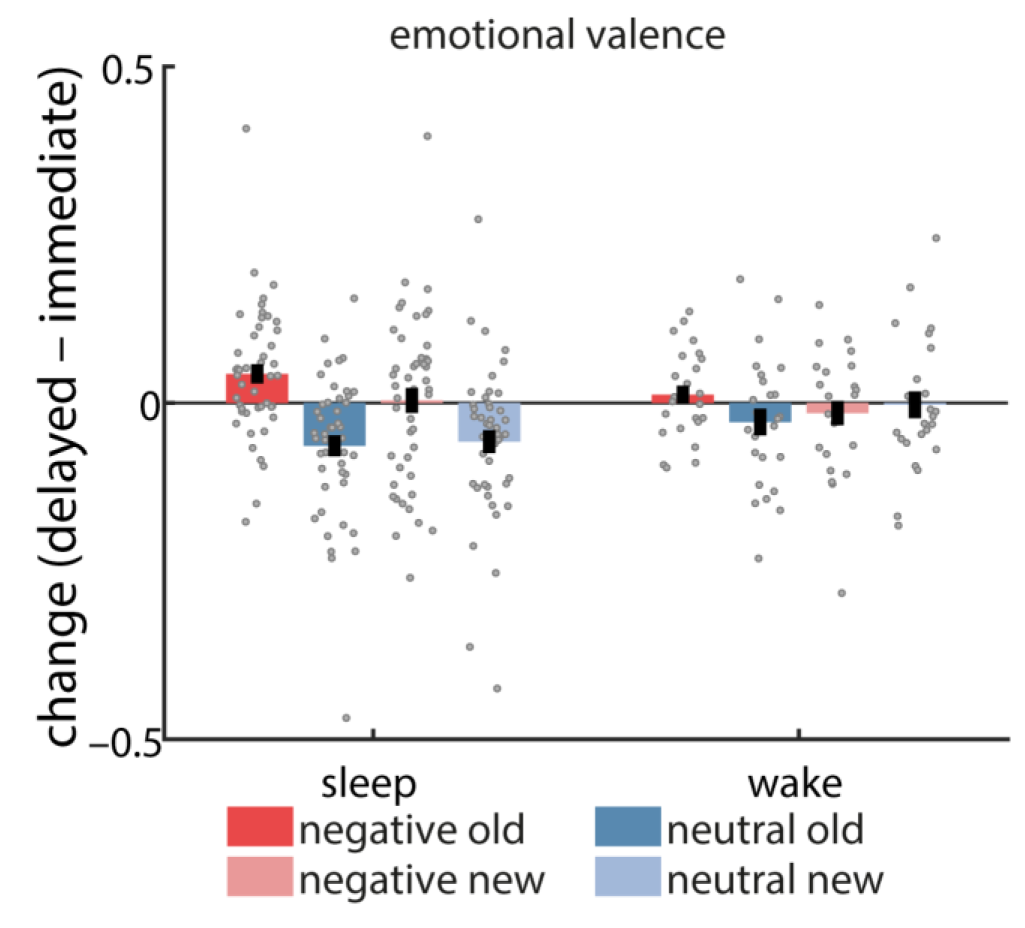
Change (delayed – immediate) in affective ratings to centrally presented items. Error bars reflect standard error of the mean with between-subject variability removed.

Specifically, sleep led to less negative ratings of negative items for old (t(45)=3.0, P=0.004), but not new (t(45)=0.2, P=0.84) items, suggesting that sleep reduces the emotionality of previously memorized negative items. Moreover, sleep led to less positive ratings of neutral items (which, overall, had positive ratings at immediate test), but did so for both old (t(45)=4.1, P=0.0002) and new (t(45)=3.4, P=0.002) items, suggesting a depotentiation of emotional reactivity to mildly positive material regardless of item novelty. Indeed, while for negative items we observed a significant old/new difference in emotional change across sleep (t(45)=3.3, P=0.002), this effect was absent for neutral items across sleep (t(45)=0.5, P=0.64). No significant old/new differences were found for wake (negative: t(24)=1.6, P=0.11; neutral: t(24)=1.8, P=0.09), although, numerically, values followed the pattern seen in sleep. In line with this observation, emotional attenuation was stronger in sleep vs. wake only for the neutral new items (t(69)=2.0, P=0.05). In contrast, sleep and wake did not differ significantly for any of the other conditions (all t(69)<1.5, P>0.16), indicating that sleep’s effect on emotional ratings was primarily the reduction of positive affect to previously unseen neutral items.

Correlational analyses with sleep parameters did not reveal associations with changes in affective ratings for any of the combinations of old/new and negative/neutral items (all P_adj_>0.94). We also examined whether valence changes were associated with the amount of forgetting of contextual memories. However, separate analyses for sleep/wake and negative/neutral revealed no significant associations, whether considering valence changes across old and new items, old items only, or correctly recognized old items (i.e., hits) only (all P_adj_>0.55).

## Discussion

While the notion that sleep is an important factor in emotional memory processing is widely accepted, little is understood about precisely how these factors interact ^39^. We found that sleep selectively stabilizes contextual aspects of negative memories, in a way not seen for neutral material, for either negative or neutral material across wake, or for recognition memory in any condition. Moreover, the proportion of time spent in NREM sleep predicted contextual memory retention in a valence-independent manner. In addition, an interval of sleep decreased affective responses to previously seen negative stimuli and to both old and new mildly positive stimuli.

These findings suggest a sleep-mediated benefit for emotional memory stabilization, but only for the spatial, contextual aspect of negative memories. The general observation of a sleep benefit for contextual but not recognition memory fits with evidence that paradigms probing associative memory are generally more sensitive to sleep effects compared to recognition paradigms ^11,12^. Studies investigating recognition have found stronger sleep-dependent effects for recollection-based recognition, which is thought to require stronger hippocampal involvement, than familiarity-based recognition ^46,47^. Combined, these findings suggest that sleep’s effect on declarative memory operates primarily on memories that rely on the binding machinery of the hippocampus ^13^.

Considering the emotional dimension, the selective sleep benefit for negative contextual memory is consistent with sleep’s negative vs. neutral advantages for associative memories of temporal order ^33^, and spatial location ^32^, although these split-night studies found this effect to depend on opposing sleep stages (NREM-rich early vs. REM-rich late sleep, respectively). More broadly, these observations fit with the notion that the sleeping brain preferentially processes information deemed of future relevance^7^. Specifically, there could be evolutionary advantages to remembering the spatiotemporal context in which a negative stimulus was originally encountered, allowing the organism to avoid such experiences in the future. The absence of a sleep benefit for neutral contextual information in our study could also be related to this prioritization process, in that preferential processing of emotional experiences could leave limited resources for the consolidation of neutral memories, although consolidation of neutral memories could still occur when only neutral items are learned (e.g., ^11,12,48,49^).

Regarding the contributions of specific sleep stages to contextual memory consolidation, we found that greater proportions of NREM (and correspondingly, lower proportions of REM) sleep are associated with improved retention. While these observations contrast with some evidence of REM sleep being primarily related to emotional memory processing ^16,17^, they add to an accumulating body of findings emphasizing the importance of NREM and its electrophysiological markers in the consolidation of both neutral and emotional memories ^20,26,28,33^. Indeed, the NREM-memory association was only observed when we pooled across emotional categories, not when we considered negative and neutral items separately. The absence of valence-specific relations may be due to the fact that sleep architectural parameters, as rather crude measures of brain state, do not have the required sensitivity to detect category-specific consolidation processes, which are likely reflected by subtler patterns of electrophysiological activity ^50–52^. Indeed, our experiment was designed to allow future examinations of such category-specific activity patterns in the sleeping brain (see below).

Overall, studies investigating the link between sleep and emotional associative memory have been far from consistent in their findings. For example, a study employing an object-face paired associates approach found reduced forgetting of neutral-neutral pairs across a nap vs. wake, but no difference for negative-neutral pairs ^34^, whereas a picture-sound paradigm reported similar sleep benefits for emotional and neutral pairs ^18^. Given that these studies focused on pairs of items, one possibility is that sleep’s effects depend on whether the type of association tested relates to links with background contextual components (in which case sleep may favor the stabilization of links with negative memories), or links between individual items (in which case the emotional advantage may be suppressed or even reversed). Complicating this view, the aforementioned split-night study that found superior picture-location memory for negative vs. neutral items across late REM-rich sleep observed the opposite pattern across early NREM-rich sleep ^32^. Another split-night study observed a similar early-night benefit for neutral picture-color associations, but did not observe this finding for picture-location associations (for which no differential negative/neutral or early/late sleep effects were found) ^25^. However, given that these studies focused on early/late night contrasts, it is unclear how specific the reported effects are to sleep. Lastly, a series of studies has reported that sleep enhances memory for central negative objects at the expense of their neutral background, with no such effect for neutral object-background pairings ^14,53,54^, which has been taken to imply that sleep weakens rather than preserves negative item-context links. However, as this paradigm tests recognition memory for individual objects and backgrounds, as opposed to directly probing the associative link between them, it is presently unclear how these and our findings relate.

In the current study, we found no sleep/wake effect on recognition memory, or any relationship between recognition memory and sleep architecture. As noted above, this is consistent with several studies reporting no effect of sleep on recognition memory ^11,12^ or familiarity-based recognition ^46,47^. However, this contrasts with reports showing enhanced overall ^19,29^ or negative ^15,17,23–25^ item recognition after sleep, in some cases relating these effects to REM sleep ^17, 23–25^. Given the many differences in experimental design between these studies and ours, it remains to be seen under what conditions recognition metrics are sensitive to sleep-mediated consolidation processes.

A prominent theoretical account posits that sleep, and REM sleep in particular, serves to preserve the content of emotional memories while simultaneously removing their affective layer ^55^. While the “emotional blunting” aspect of this model has received physiological support ^35,36^, subjective ratings of arousal and/or valence have found inconsistent sleep-related modulation of emotional tone ^18,19,23,24,32,37,38^, as reviewed in the introduction. Our findings provide some support for a role of sleep in reducing the emotional tone of memories, in that sleep reduced the perceived emotionality of both negative and mildly positive stimuli. For negative items, this effect was specific to previously seen items, suggesting a sleep-mediated emotional depotentiation of existing negative memories. In contrast, the decrease in emotional intensity for mildly positive pictures was similar for old and new items, suggesting that this overnight reduction in emotional reactivity was not specific to previously encoded materials. However, these results should be interpreted with care as emotional attenuation was significantly stronger across sleep than wake only for new neutral items. Moreover, changes in emotional ratings were unrelated to changes in contextual memory, or to aspects of sleep macrostructure, providing no direct support that sleep-mediated reductions in emotional reactivity go hand in hand with memory consolidation or occur in a particular sleep stage.

While the current paper reports behavioral evidence that sleep mediates the consolidation of contextual, negative memories, our study was designed to link these behavioral findings to neural evidence of category-specific memory reactivation. Specifically, we implemented the valence-VF manipulation at encoding to differentially “tag” the two cerebral hemispheres with emotional material, leading to the prediction that the hemisphere encoding negative items is more strongly activated during sleep. To achieve this goal of differential tagging, negative and neutral items were presented exclusively in one VF. Consequent to this design choice, the link between VF and emotional valence was apparent to the majority of subjects, presenting a possible confound. However, control analyses at immediate test indicated that while item valence and side ratings were not independent, this relation differed for old and new items, and contextual performance was better for hits than for misses or correct rejections, suggesting that the selective sleep benefit for negative contextual memories reflects a genuine memory effect.

Unexpectedly, we did not observe a baseline difference in side placement between hits and false alarms for negative items. In addition, the factor SIDE (negative-left/negative-right) affected contextual memory performance (Supplementary Information), although these effects were not in line with the notion of preferential processing of negative information by the right hemisphere ^40–42^. While we deem it highly unlikely that the entirety of our primary findings is due to these factors, future studies should attempt to tease apart their contributions.

In conclusion, we have provided evidence that sleep selectively stabilizes contextual aspects of negative episodic memories, possibly during NREM sleep, and may aid in attenuating the affective response to both negative and mildly positive materials. These findings lay the foundation for more detailed investigations into the electrophysiological underpinnings of sleep-related reactivation processes for emotional and contextual information. Nonetheless, the determinants of emotional processing during sleep remain enigmatic, and continued effort will be needed to unravel the complex and important interrelations among sleep, memory, and emotion.

## Methods

### Participants

Seventy-one young, healthy participants from the Boston area with normal or corrected-to-normal vision (24 male, 47 female; mean age ± SD: 21.3 ± 3.1, range: 18–33) were included in this study. All told, 68/71 participants reported being right-handed (one reported being ambidextrous, handedness data for two subjects were lost), and none reported any history of neurological, psychiatric, or sleep disorders (except for one subject reporting a past anxiety disorder). Sleep logs indicated an average of ~8 h of sleep across the three nights preceding the study (477 ± 51 min; range: 350–505). Participants were instructed to refrain from consuming recreational drugs or alcohol in the 24 h prior to the study, and to consume no more than one caffeinated beverage on the day of the study. They were instructed not to nap on the day of the first session (for the sleep group) or in between sessions (for the wake group). Post hoc evaluation of study questionnaires indicated that three subjects from the sleep group had not complied with this instruction. However, given that overnight sleep parameters appeared normal (total sleep time > 425 min; sleep efficiency > 78%), they were included. Two additional participants were excluded because their valence ratings at encoding and/or testing failed to differentiate between negative and neutral items, suggesting poor compliance with instructions. Subjects were compensated monetarily for their participation. All subjects provided written informed consent, the experiment was carried out in accordance with relevant guidelines and regulations, and this study was approved by the institutional review board of Beth Israel Deaconess Medical Center.

### Protocol Overview

The protocol (Fig. 1A) consisted of two sessions separated by ^~^12 h containing either sleep or wakefulness (between-subject factor GROUP). The primary components of Session 1 were encoding followed by an immediate test (Test 1), whereas the primary component of Session 2 was a delayed test (Test 2; see below for detailed descriptions of each session). Participants in the sleep group (N=46) completed Session 1 in the evening, spent the night in the sleep lab, and completed Session 2 the next morning, whereas wake subjects (N=25) came to the lab for separate morning and evening visits. An additional between-subject factor SIDE defined the association between VF and emotional category (negative-left/neutral-right: N=36 [sleep: 22; wake: 14]; negative-right/neutral-left: N=35 [sleep: 24; wake: 11]). Males and females were equally distributed across GROUP and SIDE (χ^2^=0.79, P=0.94).

Upon arriving at the lab, subjects in the sleep group were wired for high-density EEG recordings. Wake participants were wired with a simple electrooculography (EOG) montage in order to have similar setup procedures, experiences, and compliance with instructions in both groups. Session 1 typically started between 9–10 PM (sleep) or 9:15–10:15 AM (wake), and lasted 98 ± 13 min (range: 80–158). Whereas wake subjects left the lab between sessions, sleep participants were given a 9 h sleep opportunity (lights out between 10:30 PM–12:15 AM). Immediately following waking, sleep participants were asked to provide details regarding dream content and sleep quality, and then had breakfast. Session 2 commenced between 8:30–9:30 AM (sleep) or 9–10 PM (wake), lasting 89 ± 16 min (range: 60–149). The ~12 h interval between the start of Sessions 1 and 2 differed slightly, but consistently, between groups (sleep: 704 ± 20 min; wake: 722 ± 18 min; t(69)=3.9, P=0.0002).

Of note, approximately two-thirds of subjects (46 vs. 25) were assigned to the sleep condition. Given that the primary objective of our EEG recordings was to allow future examinations of hemisphere-specific memory reactivation during sleep (see Discussion), the sleep group required a sufficient number of subjects in each SIDE condition to enable negative-left vs. negative-right comparisons of electrophysiological activity. In contrast, while the wake group was critical for testing the sleep-dependence of behavioral effects, EEG collection and comparisons based on SIDE were not deemed central for the wake group, allowing for smaller group size.

### Stimuli

A total of 384 pictures (192 negative, 192 neutral) were selected from the Nencki Affective Picture System ^56^ (NAPS), a collection of affective pictures providing extensive normative valence and arousal ratings, semantic category labels, and detailed descriptions of physical image properties. Normative valence and arousal scores, based on N=204 raters, ranged from 1 (very negative; relaxed) to 9 (very positive; aroused), with 5 being neutral. Picture selection was limited to items of 1600 x 1200 pixels (landscape orientation) to ensure identical dimensions during presentation. Negative items were chosen that had normative ratings between 2–4.5 (valence) and 6–9 (arousal), while neutral items had ratings of 5–7 for valence and 2–5 for arousal, resulting in non-overlapping distributions for the two emotional categories. The number of negative and neutral items within each semantic category was matched (faces: 60 negative and 60 neutral; animals: 42 of each; objects: 42; people: 36, landscapes: 12). In addition, we selected negative and neutral items within each semantic category such that physical image properties were minimally different between emotional categories. Selected negative and neutral items differed significantly in terms of valence (3.3 ± 0.6 vs. 6.2 ± 0.5; t(382)=50.0, P<10^−168^) and arousal (6.3 ± 0.5 vs. 4.4 ± 0.4; t(382)=–44.6, P<10^−152^), but not in terms of luminance, contrast, complexity (as approximated by JPEG file size), or dimensions of CIELAB color space (all P>0.24).

### Task

Task presentation was controlled using Psychtoolbox for Matlab (the Mathworks, Natick, MA). For sleep subjects, stimulus presentation markers were sent to the amplifier to allow synchronization of EEG activity to events of interest. Subjects were seated in front of a 27-inch computer display with their eyes approximately 60 cm away from the center of the screen. Subjects were instructed to keep their head in a steady position and to minimize eye blinking throughout the experiment.

For each subject, affective stimuli were randomly partitioned at runtime according to the following criteria. Two-thirds of the stimuli (256: 128 negative, 128 neutral) were reserved for encoding and to serve as “old” stimuli during retrieval tests, with separate subsets used for Test 1 and Test 2 (64 negative and 64 neutral per test). The remaining third (128: 64 negative, 64 neutral) served as foils during retrieval (“new”), again using separate subsets for Test 1 and Test 2 (32 negative and 32 neutral per test). Moreover, the relative proportion of items within each semantic category was kept identical across encoding and retrieval tests, between negative and neutral items, and between old and new items.

#### Session 1 – Pre-encoding

Before EEG or EOG wiring, subjects indicated their level of vigilance on the Stanford Sleepiness Scale ^43^ (1: wide awake; 7: no longer fighting sleep) and the question “How would you describe your ability to concentrate right now?” on a visual analog scale (range: 0–100). Following wiring, Session 1 commenced with an extensive set of instructions regarding the encoding phase. Subjects practiced fixating, using the response slider, and reporting saccades (each described, as relevant, below). Subjects were not told their memory would be tested, nor were they explicitly informed of the relation between VF and stimulus emotionality.

#### Session 1 – Encoding

Encoding trials were presented in 8 blocks of 32 trials, with pauses between blocks. Subjects were encouraged to use pauses to move, stretch, and blink, in order to enable optimal compliance with task instructions during the subsequent block. Trial order was randomized with the constraint that the same emotional category could not be presented more than three times in a row. Each encoding trial began with a white fixation dot presented for an interval jittered uniformly between 1,000 and 2,000 ms (Fig. 1B). Next, with the fixation point still present, an affective stimulus was presented for 1,000 ms in either the left or the right VF (depending on emotional category and the subject’s VF-emotional category assignment). The lateralized stimulus was presented at a size of 21.8 x 16.4 cm, with the figure center displaced 13.9 cm laterally (13° of visual angle at 60 cm from the screen) and 4.2 cm downward (4°), resulting in a stimulus covering 20° x 15° of visual space. Another fixation display followed for 1,000 ms. Next, a screen appeared asking subjects to rate the perceived emotionality of the preceding image (“How EMOTIONAL would you say this picture is?”). Subjects indicated their response by placing a mouse-operated cursor on a vertically oriented continuous slider (visual analog scale) containing five equally spaced tick marks with associated labels (from bottom to top: very negative, somewhat negative, neutral/unsure, somewhat positive, very positive). Upon clicking, the slider position was recorded as a valence rating in the interval from –1 to +1, a blank display was presented for 500 ms, and the next trial commenced. No time restrictions were imposed for responses.

Subjects were instructed to keep their gaze on the fixation dot when present and to minimize eye blinks and other movements; they were free to move their eyes while the response screen was displayed. In addition, subjects were asked to click the mouse before onset of the response screen in the event that they made a saccade during the immediately preceding stimulus presentation. In that case, a confirmation (“You indicated you looked at the picture…”) was displayed for 2,000 ms, and the interrupted trial was repeated. On average, subjects reported making 12.6 ± 14.9 saccades (range: 0–65) during the 256 trials. Although it is unclear how reliably self-reported saccades reflect actual saccades, repeated instructions at the beginning of each block to report saccades, as well as saccades being reported by 65/71 subjects, likely helped maintain central fixation. To further dissuade subjects from making undesired movements, a warning (“Do not press any keyboard buttons unless you’re asked to!”) was presented for 2,000 ms upon detection of any key press, and the interrupted trial was repeated. Trials were repeated to maximize the number of trials available for future EEG analyses. However, behavioral findings were unaltered when excluding repeated trials.

Following the encoding phase, subjects used the slider (range from –1 to +1) to rate: 1) the visibility of laterally presented stimuli (ranging between “it was a complete blur” – “I saw everything perfectly”; sleep: 0.15 ± 0.34; wake: 0.22 ± 0.32), 2) difficulty fixating (“extremely easy” – “extremely difficult”; sleep: 0.05 ± 0.35; wake: 0.10 ± 0.41),3) accuracy fixating (“I looked away all the time” – “I never looked away”; sleep: 0.61 ± 0.30; wake: 0.51 ± 0.34), and 4) accuracy reporting saccades (“I was too slow or forgot to click” – “I perfectly reported every time I looked”; sleep: 0.49 ± 0.43; wake: 0.58 ± 0.41). None of these parameters differed between the sleep and wake groups (all t(69)<1.3, all P>0.22).

#### Session 1 – Immediate Retrieval (Test 1)

Subjects were informed that a surprise memory test would be administered, and received instructions. Retrieval trials were presented in 6 blocks of 32 trials, with pauses between blocks. Trial order was randomized with the constraint that the same combination of emotional category and old/new status could not be presented more than three times in a row. Retrieval trials (Fig. 1C) consisted of a sequence of pre-stimulus (1,000–2,000 ms), stimulus (1,000 ms), and post-stimulus (1,000 ms) periods. Critically, stimulus presentation was now central (as opposed to lateralized), with similar stimulus dimensions as during encoding. Following the post-stimulus fixation display, a first response screen asked subjects to indicate the preceding item’s OLD/NEW status (“How confident are you that this picture was shown during the LEARNING phase?”) using a response slider with equally spaced labels (certainly old, probably old, unsure, probably new, certainly new). A second response screen asked subjects for a continuous left/right judgment (“On what SIDE of the screen would you place this picture?”), with labels reading certainly left/probably left/unsure/probably right/certainly right. Subjects had been instructed to indicate where each picture “belongs”, and that their brain could still have processed the item’s left/right information even if they did not remember seeing it during encoding. Collecting left/right judgments regardless of old/new status and old/new response allowed us to compare these ratings as a function of recognition status. Finally, a third response screen asked for a valence rating as at encoding, after which a blank post-response display was shown for 500 ms, and the next trial began.

Following immediate retrieval, subjects were informed that a similar retrieval procedure would be administered during Session 2. This ensured that all subjects had similar test expectations, which are known to affect sleep-dependent memory consolidation ^57,58^.

#### Session 2

Immediately upon waking, subjects in the sleep group provided details regarding dream experiences ^59^ and sleep quality. Then, for both groups, assessments of vigilance were administered, followed by the delayed test (Test 2) using trial timing identical to Test 1. Importantly, all items employed in Test 2 (both old and new) were different from those in Test 1. In addition, a second delayed test (Test 3) was administered using the items previously used in Test 1 (Fig. 1A). However, contrary to the large drop in HR from Test 1 to Test 2 (Table 3), no significant decrease was observed from Test 1 to Test 3 in any condition (t-tests vs. zero: all P>0.27). This suggests a retrieval-related encoding effect such that at Test 3 subjects remembered the old/new judgment they had previously given at Test 1. Therefore, data involving Test 3 were not further analyzed. At the end of the protocol subjects completed an exit questionnaire probing their thoughts about the study’s goal and their awareness of the VF-valence association.

#### Resting states

Before and after every encoding and retrieval phase we collected data from 5 min resting states (six in total; Fig. 1A). Subjects were instructed to close their eyes and rest quietly. After 5 min, a sound indicated they could open their eyes again. Resting states were collected to allow investigation of waking EEG signatures related to sleep-dependent memory consolidation, and were also included for wake subjects to minimize protocol differences between groups.

### Data Acquisition and Preprocessing

EEG in the sleep group was collected using 62-channel EEG caps (Easycap GmbH, Herrsching, Germany) with channel positions in accordance with the 10-20 system. Additionally, two single cup electrodes were placed on the mastoid processes, two lateral to each eye for EOG, two on the chin for electromyography, one on the forehead as an (online) reference, and one on the skin over the clavicle as a ground. For the wake group, only EOG, mastoids, reference, and ground electrodes were connected. An AURA-LTM64 amplifier and TWin software were used for data acquisition (Grass Technologies, Warwick, RI, USA). Impedances were kept below 25 kΩ and data were sampled at 400 Hz with hardware high-pass and low-pass filters at 0.1 Hz and 133 Hz, respectively. Impedances were checked repeatedly throughout the protocol.

Sleep staging was performed in TWin using 6 EEG channels (F3, F4, C3, C4, O1, O2) referenced to the contralateral mastoid (bandpass filtered 0.3–35 Hz), bipolar EOG (0.3–35 Hz) and bipolar EMG (10–100 Hz), on 30 s epochs according to standard AASM criteria ^60^.

### Data Analysis and Statistics

All analyses were performed using custom Matlab scripts. Valence ratings, ranging from –1 to +1, were analyzed in continuous format. Continuous old/new confidence ratings were dichotomized as old (<0) and new (>0) to enable calculation of hits, false alarms, correct rejections, and misses, as well as d’ (using Macmillan and Kaplan ^61^ adjustments as required). Similarly, continuous left/right ratings were converted into (continuous) incorrect side/correct side judgments and dichotomized as correct or incorrect. Percentage of items assigned to the correct side was calculated separately for each of the aforementioned recognition classes. Items with continuous recognition or side scores of exactly zero were excluded from dichotomization. We also performed memory analyses using continuous scores, but as the pattern of results was highly similar to that obtained using dichotomized scores, results based on continuous data are not reported.

Group comparisons were performed using conventional t tests (or Wilcoxon Rank Sum for non-normal distributions) and ANOVAs with between-and within-subject factors as required. Correlations were performed using robust regression procedures to minimize the influence of outliers. In addition, Pearson or Spearman correlations (as appropriate) were performed to obtain standard estimates of correlation strength (which are not provided by robust regression). Direct comparison of correlations was performed using the procedure outlined by Meng and colleagues ^62^. Correction for multiple comparisons was done using the False Discovery Rate procedure 44 as appropriate.

## Data Availability

All data underlying the current report are available in Microsoft Excel and Matlab format (Supplementary Dataset 1).

## Acknowledgments

This work was supported by grants from The Netherlands Organization for Scientific Research (NWO) to RC (446-14-009); National Institutes of Health to ACS (F32-NS093901), and RS (MH048832, MH092638); The Harvard Clinical and Translational Science Center (TR001102); and Stanley Center for Psychiatric Research at Broad Institute.

## Author Contributions

Conceived and designed the experiment: RC, MLVB, ACS, RS. Collected data: RC, MLVB, HG. Preprocessed data: MLVB, MB, EC, EP, OPM. Analyzed data: RC, MLVB, ACS, RS. Prepared figures: RC. Wrote the manuscript: RC. All authors contributed to interpretation of results and reviewed the manuscript.

## Competing Interests

The authors declare no competing interests.

